# Microbial degradation of a widely used model polyethylene is restricted to medium- and long-chain alkanes and their oxidized derivatives

**DOI:** 10.1101/2025.09.22.675545

**Authors:** Ronja Marlonsdotter Sandholm, Gordon Jacob Boehlich, Ørjan Dahl, Ravindra R. Chowreddy, Anton Stepnov, Gustav Vaaje-Kolstad, Sabina Leanti La Rosa

## Abstract

Plastics are widely used materials, yet their chemical stability hinders biodegradation, exacerbating pollution on a global scale. Soils contaminated with plastic may foster microbes adapted to degrade plastics or plastic derivatives, and these organisms and their enzymes offer promising avenues for the development of biotechnological recycling strategies. Here, two microbial communities originating from soil collected at a plastic-contaminated landfill in Norway were enriched to select for bacteria involved in the decomposition of a commonly used, model polyethylene (PE; weight average molecular weight (M_w_) ∼4000 g/mol). We leveraged genome-resolved metatranscriptomics to identify active population affiliated with *Acinetobacter guillouiae* and *Pseudomonas* sp., showing a suite of upregulated genes (including those encoding alkane 1-monooxygenases, flavin-containing monooxygenases FMOs, cytochrome P450 monooxygenases) with functions compatible with degradation of oxidized products as well as medium- and long-chain hydrocarbons. Strikingly, spectroscopic, spectrometric and chromatographic analyses revealed the unexpected presence of medium- and long-chain alkanes and 2-ketones in the model PE substrate, preventing the erroneous conclusion that the community was interacting with the polymeric component. Consistently, only alkanes and 2-ketones with chain length of 10-35 were selectively degraded by an *A. guillouiae* isolate, as confirmed by proteomics analyses and substrate characterization following bacterial growth. Besides extending the knowledge on the enzymatic basis for degradation of PE-derivatives in soil-associated microbial systems, our results provide an advanced compositional characterization of a widely used model “PE” material, while offering valuable insight to support future studies aimed at unequivocally identifying organisms and their enzymes implicated in PE transformation.

## Introduction

Plastics are synthetic materials composed primarily of long-chain polymers derived from petrochemicals. Plastics have become integral to everyday life due to their light weight, durability, electrical insulation, moldability, and cost-effective manufacturing [1]. As a consequence, global production has surged from 464 million tons per year in 2020 to a projected 804 million tons per year by 2050 [2]. Polyethylene (PE) is the most extensively manufactured plastic, constituting approximately 36% of global plastic production [3]. It is a nonpolar, partially crystalline polymer characterized by its high chemical resistance, low density, and excellent electrical insulation. PE is commonly classified as low-density polyethylene (LDPE), known for its flexibility and transparency, and high-density polyethylene (HDPE), being more rigid and dense [4].

PE-based plastics produced worldwide are either recycled, incinerated or sent to landfills. These methods are largely inefficient and energy intensive, with landfilling in particular leading to accumulation of plastic materials with impact on the surrounding environment [5,6]. In addition, the mechanical recycling of PE decreases the quality of the material due to thermo-oxidation, crosslinking and chain functionalization [7]. In the absence of efficient mechano-chemical processes that facilitate reuse of PE, there is an urgent need to explore biological approaches capable of efficiently degrading or transforming plastic waste, thereby enhancing recycling and mitigating its environmental accumulation and impact. PE-based plastics dispersed in landfill and forest soils are subjected to prolonged exposure to molecular oxygen, water, sunlight and UV radiation. These factors can trigger chemical reactions leading to the introduction of polar functional groups (e.g., carbonyl and hydroxyl) onto the polymer backbone, thus rendering PE more accessible to depolymerization, providing a unique setting for microbial adaptation [8]. Therefore, samples from landfills and soil originating from plastic-polluted areas may serve as rich reservoirs of bacteria that have evolved the ability to exploit this novel nutrient niche [9].

To address the plastic pollution challenge, plastic-degrading microorganisms and their enzymes offer promising avenues for the development of biotechnological recycling strategies. In recent years, a multitude of studies have indicated that bacteria belonging to the genera *Acinetobacter*, *Pseudomonas*, *Rhodococcus*, *Brevibacillus*, *Aneurinibacillus* and *Bacillus*, among others [10–17] possess capabilities to depolymerize PE through different enzyme systems. These potential enzymes belong to various oxidoreductase families, such as alkane hydroxylases, laccases, manganese peroxidases and monooxygenases [9]. Despite these claims, the mechanism by which PE can be degraded by microbes in natural environments remains controversial [18]. Indeed, interpretation of data on the depolymerization and metabolization of PE is often complicated by the lack of appropriate controls, use of unspecific analytical techniques (e.g., microscopy or infrared spectroscopy), and by the reliance on non-characterized model plastics, which may contain metabolizable additives and oligomers [19].

In this study, we used a multi-omics approach to functionally characterize two microbial communities enriched on a low molar mass polyethylene-like wax (PELW), a commonly used model substrate in PE studies. The microbial communities were sourced from a plastic-contaminated landfill in Norway, and therefore they faced a prolonged exposure to plastic, including PE, prior to the experiment. We report the isolation of a bacterium, *A. guillouiae* FS11 able to grow on PELW, and employed proteomics to identify enzymes involved in the depolymerization of components derived from the PELW. Furthermore, a combined analytical approach showed that the PELW was substantially oxidized “out of the box”, a property not noted by the manufacturer, with consequence on its microbial accessibility. Besides extending the knowledge on the enzymatic basis for degradation of PE-derivatives in soil-associated microbial systems, our findings also offer a cautionary perspective on the increasing number of studies claiming enzymatic degradation of materials with insufficient compositional characterization.

## Material and methods

### Sample collection and microbial profiling using 16S rRNA amplicon sequencing

Soil was collected on the 1^st^ of November 2022 from a plastic contaminated landfill in Finnskogen (Flisa, Norway). under a permit of the regional authority of Flisa Municipality. At sampling, large amounts of macroplastic items were present on the surface and buried, having accumulated at this site since the 1970s-1980s without removal. Two locations were sampled: S1 (∼5 cm topsoil; 60°40’32.32’’N, 12°6’18.98’’E) and S2 (∼15 cm subsoil; 60°40’32.71’’N, 12°6’19.07’’E). The original S1 and S2 samples were sent to DNASense ApS (Aalborg, Denmark) for 16S rRNA amplicon sequencing. Detailed information for DNA extraction, library preparation, sequencing and data analysis is provided in the Supplementary Material.

### Substrates and growth conditions

PE powder from Sigma-Aldrich (cat. number 427772; weight average molecular weight (M_w_) ∼4000 g/mol, number average molecular weight (M_n_) ∼1700 g/mol), was used as a model commercial substrate. Because its M_w_ is far below that of conventional polyethylene (> 100,000 g/mol), we designate the material as a polyethylene-like wax (PELW) to distinguish it from high-M_w_, “common” polyethylene. Triacontane (C30) and tetracontane (C40) were obtained from Sigma-Aldrich (cat. number 263842 and 87087). D-glucose and sodium acetate were purchased from VWR (cat. number 0188-2.5KG) and Sigma-Aldrich (cat. number S2378), respectively. LDPE granules from Borealis (grade FT5230) milled to powder with 200-300 µm particle diameter. Unless stated otherwise, cultures were grown using M9 minimal salts medium (MM) supplemented with a single carbon source [20]. Microbial growth was determined spectrophotochemically by measuring optical density at 600 nm (OD600).

### Selective enrichment of soil samples on PELW and alkanes

The sequential enrichment protocol was initiated by inoculating MM containing 10 mg/mL PELW as a sole carbon source with soil from S1 and S2. These primary cultures were incubated for 10 days, then 100 µL were transferred to 20 mL of fresh MMwith PELW to obtain secondary cultures. Tertiary cultures were prepared by inoculating 100 µL of a secondary culture into three separate conditions: 20 mL of fresh MM containing PELW (resulting in the cultures indicated as S1PELW and S2PELW), 20 mL of MM supplemented with 10 mg/mL of C30 (S1C30 and S2C30), and 20 mL of MM supplemented with 10 mg/mL of C40 (S1C40 and S2C40). Incubation conditions in all enrichment phases were 30°C with 200 rpm agitation. At each enrichment step, two controls (i.e., MM without substrate or MM without any microbial source) were also set up and subjected to the same conditions.

### Shotgun sequencing and assembly of metagenomes

Samples from the tertiary communities S1PELW, S2PELW, S1C30, S2C30, S1C40 and S2C40 were collected in the early stationary phase (**Fig. S2**) by pelleting at 15,000 G for 10 min. Pellets were resuspended in 800 µL Solution CD1 from the DNeasy^®^ PowerSoil^®^ Pro kit (QIAGEN, ID: 47014) and DNA was extracted according to the manufacturer’s instructions. DNA size was selected using a Short Read Eliminator XS kit (PacBio, PN: 102-208-200). DNA was quantified using Qubit™ dsDNA Broad Range Assay (Invitrogen) on a Qubit™ 1.0 fluorometer. DNA size was assessed using gel electrophoresis, and purity was evaluated using a NanoDrop™ One Microvolume UV-Vis spectrophotometer (ThermoFisher Scientific). Sequencing libraries were prepared using the Native Barcoding kit SQK-NBD114.24 (Oxford Nanopore Technologies, UK) following the manufacturer’s instructions. S1PELW and S2PELW libraries were pooled, as were S1C30 with S2C30, and S1C40 with S2C40. Each pooled sample was then sequenced on a separate R10.4 flow cell on a MinION device (Oxford Nanopore Technologies, UK). POD5 files were base-called and demultiplexed with Dorado v0.5.0 [21], using the super-accurate model (config: dna_r10.4.1_e8.2_400bps_sup@v4.3.0).

The snakemake wrapper Aviary v0.9.1 was used to recover MAGs [22]. Low-quality MAGs, defined according to the MiMAG standards [23] were removed from the dataset. MAGs were taxonomically classified using GTDB-Tk [24] with the GTDB database release 220. Functional annotation of the bacterial genomes was performed with DRAM v1.4.6 [25] with the following databases: Uniref90, PFAM-A, KOfam and dbCAN-V10 (all downloaded in January 2024). Figures were created using R v4.3.3 [26] and ggplot2 v3.5.1 [27].

### Metatranscriptomic analysis

Samples from the tertiary communities S1PELW, S2PELW, S1C30, S2C30, S1C40 and S2C40 were collected by pelleting at 15,000 G at 4 °C in mid-exponential phase. Pelleted samples were shipped to Novogene (Cambridge, UK) for extraction and sequencing of metaRNA. Detailed information for RNA extraction and sequencing is provided in the Supplementary Material. Raw metatranscriptomic reads were quality checked using FastQC v0.12.1 [28] filtered using fastp v0.23.4 (`-q 20`) [29] and ribosomal RNA removed from the dataset using SortMeRNA v4.3.6 [30]. The filtered metatranscriptomic dataset was pseudo-aligned against the recovered MAGs using Kallisto v0.48.0 [31]. Transcripts per million (TPM) values were log_2_-transformed. Genes with transcripts in 2 out of 3 replicates were kept. The remaining missing values were imputed from a normal distribution (width of 0.3 and down shifted 1.8 standard deviations from the original distribution). To find differentially expressed genes, a paired t-test (*p* ≤ 0.05) was preformed using Rstatix v0.7.2 comparing the data from C30, C40 or PELW to a glucose control [32]. Figures were generated using R v4.3.3 [26] packages ggplot2 v3.5.1, cowplot v1.1.3 [33] and ggrepel v0.9.5 [34].

### Isolation of *Acinetobacter guillouiae* FS11 and proteomic analysis

*Acinetobacter guillouiae* was isolated from the S1PELW tertiary community following selection on MM containing 10 mg/mL PELW and Luria Bertani (LB, Sigma-Aldrich, L3522) plates supplemented with 50 µg/mL kanamycin sulfate (Gibco, 11815-032). Detailed information for the identification of *A. guillouiae* FS11 is provided in the Supplementary Material. To obtain samples for proteomic analysis, *A. guillouiae* FS11 was cultured in triplicates in MM supplemented with either 10 mg/mL PELW, 10 mg/mL C30 or 2% v/v sodium acetate. Three samples types were collected in the mid-exponential phase (**Fig. S4**): planktonic cells, surface biofilm attached to PELW, and secreted proteins. Detailed information for protein extraction, trypsin digestion, and mass spectrometry analysis is provided in the Supplementary Material.

Analysis of mass spectrometry data was done using FragPipe v21.1 (https://fragpipe.nesvilab.org/), with MSFragger v4.0 [35], IonQuant v1.10.12 [36], Philosopher v5.1.0 [37]. A sample specific database consisting of the *A. guillouiae* FS11 proteome (4615 proteins) was used. The final database was supplemented with common contaminants (e.g., human keratin, bovine serum albumin and casein) and contained reverse entries of all library sequences. Carbamidomethylation was used as a fixed modification, while oxidation of methionine and protein N-terminal acetylation were added as variable modifications. Trypsin was used as a digestive enzyme, with maximum one missed cleavage allowed. For Label-Free-Quantification (LFQ), IonQuant was applied with FDR-controlled match-between-runs (MBR) enabled. The LFQ intensity values were log-transformed, and a protein was considered “present” if it was detected in at least two of the three biological replicates in at least one condition. Missing values were imputed from a normal distribution. Differential abundance analysis was performed using a paired t-test (*p* ≤ 0.05) in Rstatix v0.7.2 [32] in R v4.3.3 [26]. Figures were generated using the packages ggplot2 v3.5.1 [38], cowplot v1.1.3 [33] and R v4.3.3 [26].

### Protein structure prediction

3D structural models of the top 50 most abundant proteins in the *A. guillouiae* FS11 PELW-proteome were generated using AlphaFold3 [39]. Structure similarity searches were performed using FoldSeek [40]. Molecular images, with computation of surface potential, were generated using UCSF ChimeraX v1.10 [41].

### Microscopy

Experimental samples consisted of 10 mg/mL PELW derived from MM inoculated with S1PELW or S2PELW soil communities, or *A. guillouiae* FS11, while control samples consisted of 10 mg/mL PELW derived from MM without any microbial source. Scanning electron microscopy (SEM) analysis was conducted using a Hitachi SU5000 microscope (Hitachi High-Technologies Corporation, Japan). The specimens for SEM analysis were prepared by spreading powdered PELW samples on an EM-Tec CT12 conductive double sided adhesive carbon tab (Delta Microscopies, France) and by sputter coating with Platinum with Cressington 208 HR sputter coater (Cressington Scientific Instruments, England). For sputter coating, 20 mA sputter current for 30 seconds and sample holder with tilt and rotation was utilized. The Platinum coated specimens were investigated for SEM under high vacuum mode with SE detector at an accelerated voltage between 1.5 – 3 kV and working distance about 10 mm.

### Substrate characterization

#### Fourier-transform infrared spectroscopy

The FT-IR spectrum of the PELW and pure LDPE (FT5230, Borealis, Austria) was acquired with a Spectrum Two FT-IR spectrometer (PerkinElmer, Waltham, MA, USA) equipped with a Quest ATR (attenuated total reflectance) sampling module (Specac Ltd, Orpington, United Kingdom). The signals were obtained with 4 cm^-1^ spectral resolution (8 consecutive readings per measurement) in the 4000-550 cm^-1^ range using Spectrum IR software (PerkinElmer, Waltham, MA, USA). The carbonyl index was determined by calculating the ratio of the absorbance peak at 1715 cm⁻¹ to the reference peak at 1505 cm⁻¹.

### Pyrolysis-gas chromatography-mass spectrometry

Samples consisting of 150-250 µg of PELW or a commercial LDPE (grade 22H594; INEOS Olefins Switzerland), were placed in a deactivated stainless-steel sample cup and introduced into a multi-shot EGA/PY-3030D pyrolyzer (Frontier Laboratories, Japan) coupled to a 7890N gas chromatograph and a 5975 mass selective detector (Agilent Technologies, USA). The furnace temperature was maintained at 550 °C and the interface between the furnace and the GC-MS system was set to 300 °C. The GC injector was operated in split mode (100:1 ratio) at 300 °C. The analysis of volatile products was performed using an Ultra-Alloy metal capillary column containing 5 % diphenyl- and 95 % dimethylpolysiloxane stationary phase (30 m, 0.25 mm ID, 0.25 µm; Frontier Laboratories Ltd., Japan) using helium as carrier gas at 1 ml/min flow rate. The GC oven temperature increased from 70 °C to 350 °C at 20 °C/min and maintained at 350 °C for 16 min. The electron ionization (70 eV) mass selective detector was operated with an ion source temperature of 230 °C, interface temperature of 300 °C, and a scan range of 29-350 m/z.

Compound identifications were done by matching chromatograms using an F-Search in the Wiley 11^th^ edition/NIST 2017 spectral library. To refine identification, spectral match scores were combined with retention times to differentiate between compounds with similar spectra.

### ^1^H and ^13^C nuclear magnetic resonance spectroscopy

^1^H-NMR and ^13^C-NMR spectra of PELW and PE wax samples were recorded at 328 K using a Bruker AVANCE III HD 400 MHz instrument equipped with a BBFO room temperature probe. For sample preparation, 5 mg of pristine PELW and a PE wax (Licocene 5301; Clariant AG, Switzerland) were suspended in 0.5 mL CDCl_3_ and transferred to a 5 mm Norell^®^ Sample Vault Series™ NMR tube. Heating these samples to 328 K gave clear solutions. Chemical shifts are reported in ppm with the solvent signals (7.26 ppm for ^1^H/ 77.16 ppm for ^13^C) as internal standard. The following pulse programs where used: “`zg30`” for ^1^H, “`deptqgpsp.2`” for DEPTQ-^13^C, “`hsqcetgpsi2`” for HSQC and “`hmbcetgpl3nd`” for HMBC Phase contrast and scanning electron microscopy.

### Size exclusion chromatography and gas chromatography-mass spectrometry

Three types of samples (untreated PELW, PELW incubated in MM without any microbial source and PELW incubated in MM with *A. guillouiae* FS11) were prepared to assess the impact of *A. guillouiae* FS11 on substrate composition for subsequent GC-MS and SEC analysis (see Supplementary Methods). The weight average molecular mass (M_w_), number average molar mass (M_n_), and the molecular weight distribution in the PELW samples were determined using a GPC-IR5 system (Polymer Char, Spain) equipped with four PLgel 20 µm MIXED-A columns (Agilent Technologies, USA). Approximately 4 mg of a pure LDPE (FT5230, Borealis, Austria), PELW and a PE wax (Licocene™ 5301; Clariant AG, Muttenz, Switzerland) were dissolved in 8 ml of 1,2,4-trichlorobenzene by mixing at 160°C for 3 hours. For SEC analysis, 200 µl of the samples were injected into the SEC system. The analysis was performed at 150 °C using a high-sensitivity infrared detector and 1,2,4-trichlorobenzene as a mobile phase (1.0 mL/min flow rate). Polystyrene standards with a narrow M_w_ distribution and M_peak_ (most abundant M_w_ in the sample) in the range of 1140 – 7,500,000 g/mol (Agilent Technologies, USA) were used for calibration.

For GC-MS, extraction of low molecular weight compounds was conducted from approximately 100 mg of PELW by mixing with 3 mL of ethyl acetate in PTFE/Silicone septum sealed glass vials. The vials were heated at 95°C for 1.5 hours and the liquid fraction filtered through a 0.2 µm Teflon syringe filter and subjected to GC-MS analysis.

GC-MS analysis was carried out using an Agilent 6890N gas chromatograph (Agilent Technologies, Inc., USA) coupled to an Agilent 5973 Network Mass Selective Detector (Agilent Technologies, USA) and a Gerstel MPS2 Autosampler (GERSTEL GmbH & Co.KG, Germany). The gas chromatograph was equipped with 30 m Zebron ZB-5MSPlus column having a 250 µm internal diameter and 0.25 µm coating thickness (Phenomenex, USA). The temperature of the column increased from 60 °C to 300 °C at a heating rate of 10 °C per minute with a total run time of 45 minutes. Helium with a purify grade of 6.0 was used as a carrier gas at a flow rate of 3 mL/minute. The mass spectrometer was operated in electron ionization mode at 70 eV. Mass spectra and the total ion chromatograms were obtained by automatic scanning a mass range (m/z) of 33-720. Three runs per sample (n=3) were performed. The volatile components were identified by comparing the mass spectrum with those available in the Wiley 11^th^ edition/NIST 2017 spectra library. Compounds were quantified by calculated response factors. Response factors were calculated by using the reference compounds BHT (butylated hydroxytoluene) and Tinuvin® 120 (2’,4’-Di-tert-butylphenyl 3,5-di-tert-butyl-4-hydroxybenzoate), as they give differences in the responses between lower and higher molecular weight components.

## Results

### Microbial communities from a plastic-enriched landfill include metabolic active bacteria that grow on PELW, but not on LDPE

As part of an effort to discover and characterize microbes and their enzymes able to degrade LDPE and PELW, soil samples were collected from a landfill that has been exposed to plastic waste since the 1970s. Samples were collected from two different locations and depths and subjected and subjected to 16S rRNA gene amplicon sequencing, which revealed species previously associated with the degradation of various plastics and alkanes (**Supplementary results and Fig. S1**).

An enrichment protocol by sequential passaging was employed in order to select microbial species specifically involved in the degradation of LDPE, PELW and two distinct alkanes (triacontane and tetracontane). The two alkanes were used as substrates since they may trigger the catabolic systems also used for PE degradation. Cultures showed growth on PELW, triacontane and tetracontane (**Fig. S2**). In contrast, no growth was observed when LDPE was provided as the sole substrate.

Long-read metagenomic sequencing of the enrichment cultures yielded a total of 84 high-quality and 46 medium-quality dereplicated MAGs (**Fig. S3**). The use of long-read DNA sequencing ensured that 127 MAGs encoded full-length 16S rRNA genes, which enabled searches for the occurrence of these bacteria in the amplicon dataset generated from the original soil samples. At a 97% identity cut-off when comparing the 16S rRNA sequences derived from the MAGs in the enrichment cultures to OTUs, 125 of 130 MAGs were detected in the original soil microbial communities S1 and S2 (**Fig. 1A**). Of the undetected MAGs, 3 of 5 MAGs were missing a 16S rRNA gene. The S1PELW and S1C30 enrichments were characterized by the dominance of MAG107 (55.81%) and MAG13003 (62.81%), respectively, both taxonomically identified as *Acinetobacter guillouiae* (**Fig. 1B, Table S1A**). *A. guillouiae* was absent from the S1C40 community, suggesting its limited capacity to metabolize alkanes with a chain length >30. The S2PELW and S1C40 consortia were characterized by a high relative abundance (42-62%) of MAGs affiliated to the genus *Pseudomonas*. Scanning electron microscopy (SEM) provided an initial qualitative insight into the distribution and morphological diversity of microbial cells with the ability to adhere to the substrate. Species with distinct morphologies were observed in the S1PELW and S2PELW communities (**Fig. 1C and 1D**). In both samples, microorganisms formed dense and slimy clusters on the PE particles, resembling biofilms. Additional 16S RNA amplicon sequencing of the bacteria attached to the PELW surface confirmed the presence of *Acinetobacter* and *Pseudomonas*, respectively, in agreement with their prevalence in the corresponding enrichment community.

**Fig. 1.**
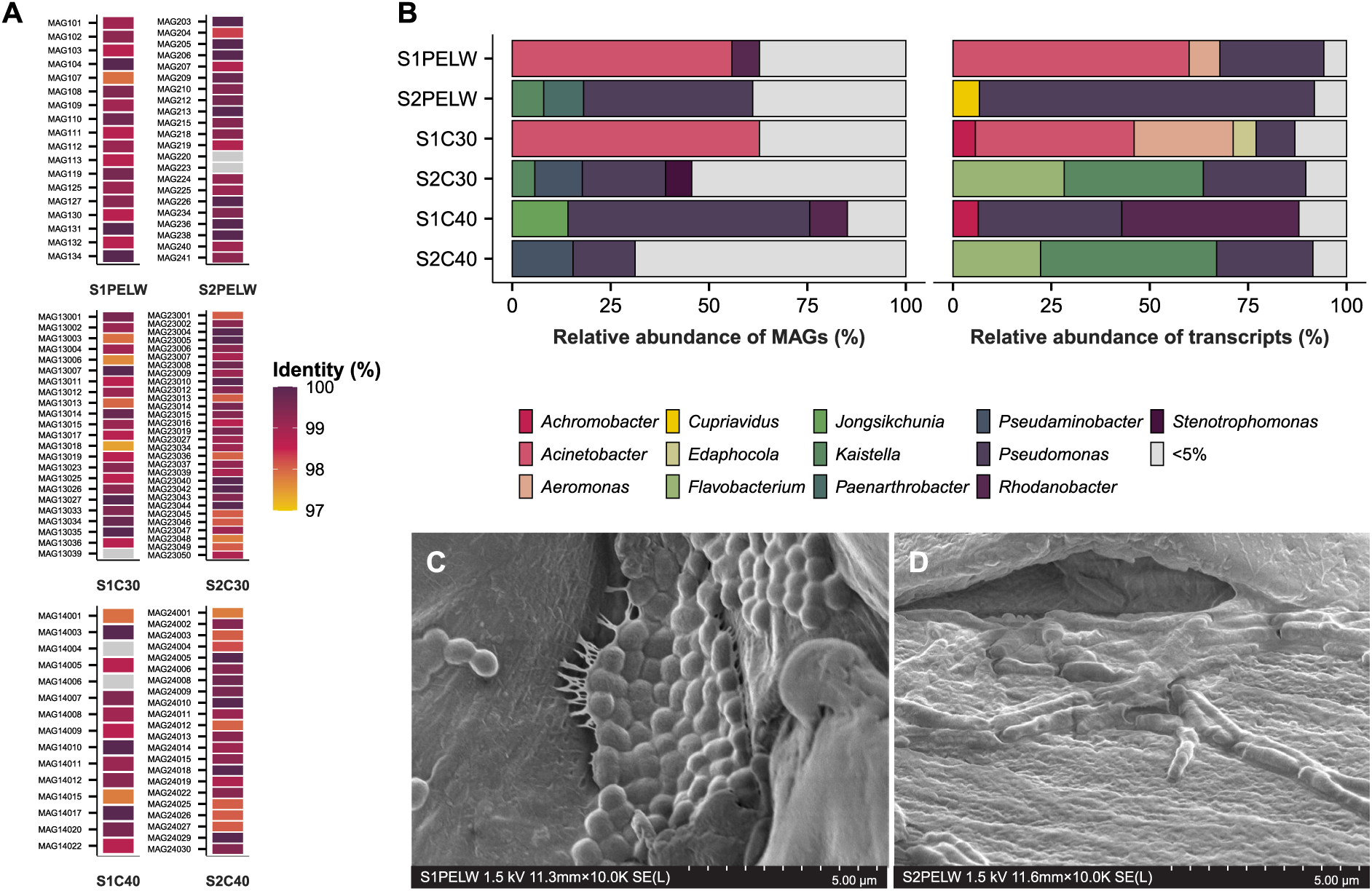
Relative abundance and activity of MAGs in the PELW, triacontane (C30) and tetracontane (C40) enrichment cultures as well as their detection in the original soil communities. **(A)** A list of high- and medium-quality MAGs reconstructed from the different enrichment cultures and their identification in the original S1 and S2 soil communities. The detection is based on the alignment of 16S rRNA gene sequence of each individual MAG to the OTUs in S1 and S2. The 16S rRNA gene detection is colored based on the % identity of the gene alignment. At a 97% identity level to OTUs, 96% of the MAGs were detected in the soil samples. **(B)** Relative abundance of MAGs (grouped at genus level) in each enrichment and the relative abundance of total transcripts per million (TPM) for each MAG within each community. Genera with a relative abundance of <5% are grouped together. Panels **(C)** and **(D)** show SEM images of the PELW surface following growth with the S1PELW and S2PELW communities, respectively. Each community exhibits a distinct morphology of bacteria attached to the PELW surface.

A genome-resolved metatranscriptomics approach allowed us to identify metabolically active MAGs and their upregulated genes in response to PELW, triacontane and tetracontane to a control where glucose was used as the sole carbon source. Intriguingly, transcripts abundance (relative to the total TPMs) for each MAG in the communities did not always reflect their abundance in the enrichments (**Fig. 1B**). In the S1PELW community, the most abundant, and most active MAG, was MAG107 (*A. guillouiae*) (**Fig. 1B**). The second most active MAGs, MAG108 (*Pseudomonas*) and MAG134 (*Aeromonas*), were not prevalent MAGs in the enrichment community (3.2% and 0.83%, respectively). As for the S2PELW community, MAG238 and MAG206 (*Pseudomonas*) were highly active, with MAG238 not being the most predominant *Pseudomonas* in the enrichment (8.33%). This observation applies to all the enrichments, where the most abundant MAGs were not necessarily the most active, highlighting the need for functional data to accurately characterize true metabolism of substrates by microbial community members that may be dormant or slow growers.

Closer examination of the metatranscriptome showed that in the S1PELW community MAG107 (*A. guillouiae*) accounted for all upregulated genes associated with the alkane terminal oxidation pathway including those coding for an alkane 1-monooxygenase (*alkM*) and the electron-transfer components rubredoxin (*rubB*) (**Fig. 2A**). In addition, several genes encoding flavin-containing monooxygenases (FMOs) were upregulated. While MAG107 appears to be the principal driver of substrate degradation, other MAGs including MAG134 (*Aeromonas aquatica*) and MAG108 (*Pseudomonas* sp.) displayed upregulated genes coding for enzymes involved in β-oxidation of fatty acids (**Table S1B**). In the S2PELW community, several MAGs exhibited upregulated genes encoding enzymes contributing to alkane degradation (**Fig. 2B**). Here, MAG238 (*Pseudomonas* sp002029345), MAG206 (*Pseudomonas fluorescens*) and MAG234 (*Pseudomonas veronii*) show increased expression of genes coding for aldehyde dehydrogenases (*aldH*) and alcohol dehydrogenases (*yiaY*, *adhP*, *ADH* and *exaA*). In this community, genes coding for cytochrome P450s (CYPs), which have been associated with both terminal and subterminal hydroxylation of alkanes [42], were upregulated in MAG238, suggesting its potential role as main degrader (**Table S1C**).

**Fig. 2.**
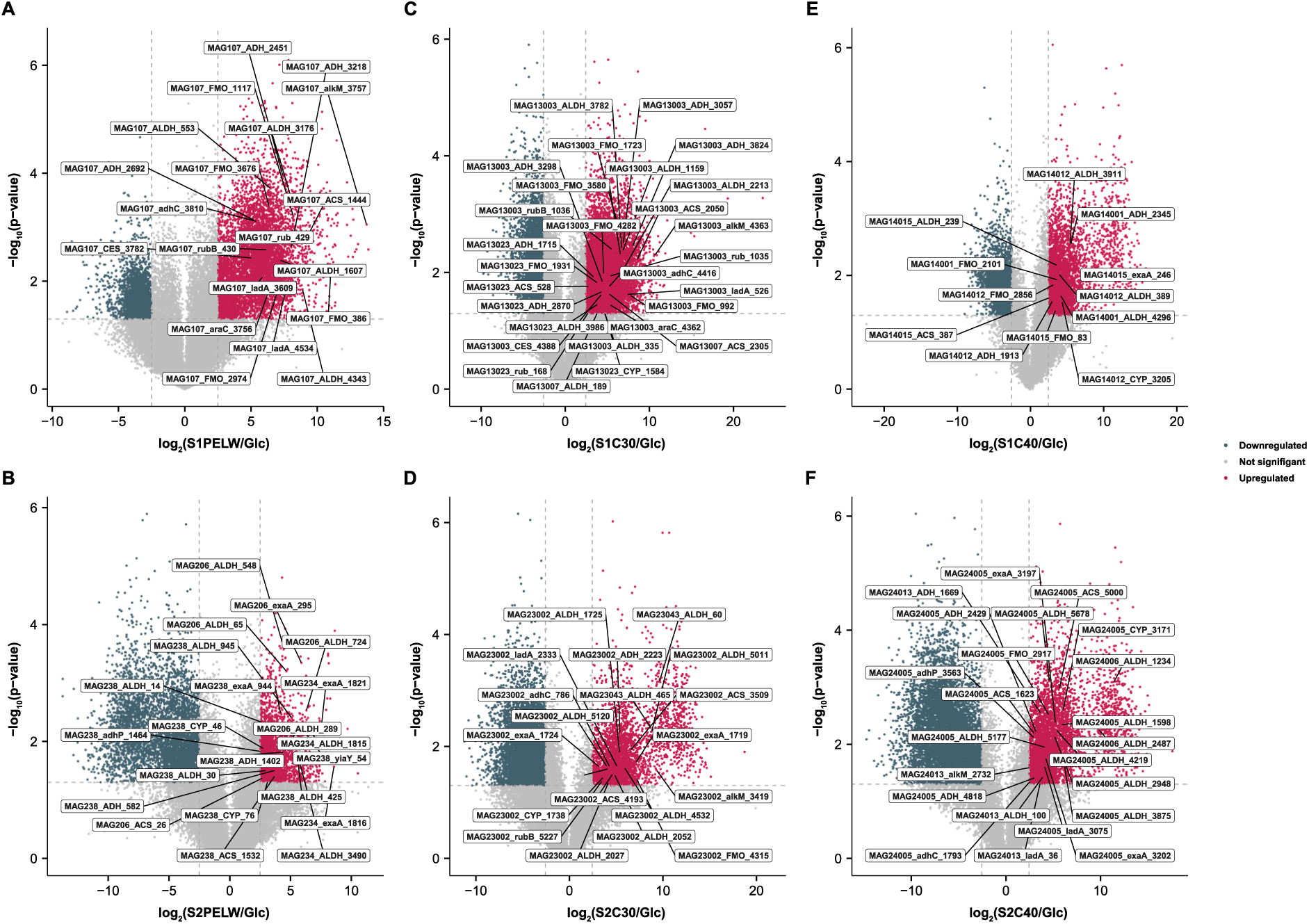
Volcano plot indicating different transcripts in the PELW, triacontane (C30) and tetracontane (C40) degrading communities that displayed a large magnitude of fold-changes in TPM and high statistical significance (-log_10_ of p-values, t-test) compared to the same communities grown on glucose. Panel **(A)** and **(B)** display upregulated genes in the S1PELW and S2PELW tertiary communities, respectively. Panel **(C)** and **(D)** show upregulated genes in the S1C30 and S2C30 tertiary communities, respectively. Panel **(E)** and **(F)** show the upregulated genes in the S1C40 and S2C40 tertiary communities, respectively. KEGG Orthology IDs were used to identify and highlight genes known to be involved in alkane degradation, 2-ketone degradation and β-oxidation of fatty acids. The dashed horizontal line denotes the p-value cut-off (*p* < 0.05), and the dashed vertical lines denote the log_2_-fold change cut-off (< −2.5 and > 2.5). Source data are provided in Table S1B-G.

The S1C30 community was dominated by MAG13003 (*A. guillouiae*), which exhibited upregulation across the full suite of genes involved in alkane degradation (**Fig. 2C**). In addition, MAG13023 (*Achromobacter* sp.) upregulated genes encoding alcohol dehydrogenases and aldehyde dehydrogenases, as well as a CYP, suggesting a potential role in alkane transformation. Upregulated genes for FMOs were detected in MAG13003, potentially oxidizing 2-ketones generated by CYP’s activity. Although MAG13023 was not one of the most active MAGs in the community (**Fig. 2C**), this MAG and MAG13003 seem to share the role as main degraders, whereas MAG13012 (*Achromobacter kerstersii*), MAG13007 (*Aeromonas aquaticia*) and MAG13006 (*Pseudomonas putida*) have genes in the β-oxidation pathway upregulated (**Table S1D**). In the S2C30 community, MAG23002 (*Pseudomonas* sp.) showed upregulated genes in the alkane degradation pathway, including *alkM*, the long-chain alkane monooxygenase *ladA* and *cyp*, as well as the β-oxidation pathway, suggesting its role as main degrader (**Fig. 2D**). MAG23043 (*Kaistella soli*) displayed upregulated genes encoding enzyme involved in the β-oxidation pathway (**Table S1E**).

In the S1C40 community, MAG14012 (*Achromobacter* sp.) upregulated genes encoding a CYP, as well as genes encoding enzymes involved in the full alkane degradation pathway and the β-oxidation pathway were detected (**Fig. 2E, Table S1F**). Other active MAGs (MAG14015; *Pseudomonas* sp, MAG14001; *A. guillouiae* and MAG14011) lacked the initial alkane hydroxylation step but showed upregulation of genes coding for enzymes involved in the subsequent alkane oxidation and β-oxidation pathway, as well as MAG14001 upregulated a gene coding for an FMO. In the S2C40 community, MAG24005 (*P. fluorescens*) and MAG24013 (*P. veronii*) showed upregulated genes coding for AlkM, LadA, with MAG24005 also upregulating a gene coding for a CYP (**Fig. 2F, Table S1G**).

Overall, comparison of the community transcriptional response in PELW versus S1C40 and S1C30 revealed no upregulated genes in PELW encoding enzymes potentially capable of cleaving the C– C bonds of the polymeric component.

### PELW (Sigma-Aldrich PE) is oxidized at the chain ends

Many studies have reported microbial degradation of the Sigma-Aldrich low molecular mass PE (i.e. catalog number 427772) [15,43–50], referred to as PELW in this study, a phenomenon reproduced by our enrichment cultures (**Fig. S2**). Since we observed no growth on pristine LDPE and no differentially expressed gene that could be linked to enzymes able to interact with the polymeric substrate, it was of interest to perform a deeper analysis of the PELW material to identify the constituents that may be responsible for sustaining microbial growth. SEC analysis showed that the PELW M_w_ (weight average molecular mass) is ∼3900 g/mol, which is similar to what is reported by the manufacturer (∼4000 g/mol). This is indeed substantially lower than a reference LDPE, with M_w_ of 97 300 g/mol, and slightly lower than a PE wax (∼5400 g/mol) (**Table S2A**). To determine the putative presence of functional groups from oxidized derivatives, PELW and LDPE were subjected to FTIR analysis (**Fig. 3B**). To our surprise, a strong peak at ≈1722 cm^-1^ was observed in the PELW sample corresponding to stretching of a carbonyl group in an aldehyde or a ketone, accompanied by additional much weaker peaks at ≈1409 and ≈1158 cm^-1^. The LDPE sample showed no sign of oxidation. To identify the type and extent of oxidation of the PELW, the polymer was analyzed by NMR. To determine the presence of ketones in the PELW, and to determine the positions of the carbonyl groups, the PELW structure was elucidated using one- and two-dimensional ^1^H and ^13^C-NMR (**Fig. 3C**). The ^1^H-NMR spectrum displayed a large singlet at 1.28 ppm, which is characteristic of the CH_2_-groups within the hydrocarbon backbone of PE. As expected, this peak was present in both the PE wax and the PELW samples. However, additional signals from the PELW were detected between 2.00-2.50 ppm. The chemical shift of these signals is typical for protons located next to carbonyls. The singlets at 2.10 and 2.12 ppm are characteristic of CH_3_ groups next to a carbonyl. The two triplets at 2.37 and 2.40 ppm indicate CH_2_ groups next to a carbonyl. The ^1^H-^13^C-HMBC-spectrum showed correlations from the triplet at 2.40 ppm and the singlet at 2.12 ppm to a carbonyl signal at 208.7 ppm, confirming that these signals stem from a methyl ketone (2-ketone). The smaller triplet at 2.37 ppm showed a correlation to a carbonyl signal at 211.3 ppm, indicating this triplet stems from in-chain ketones.

**Fig. 3.**
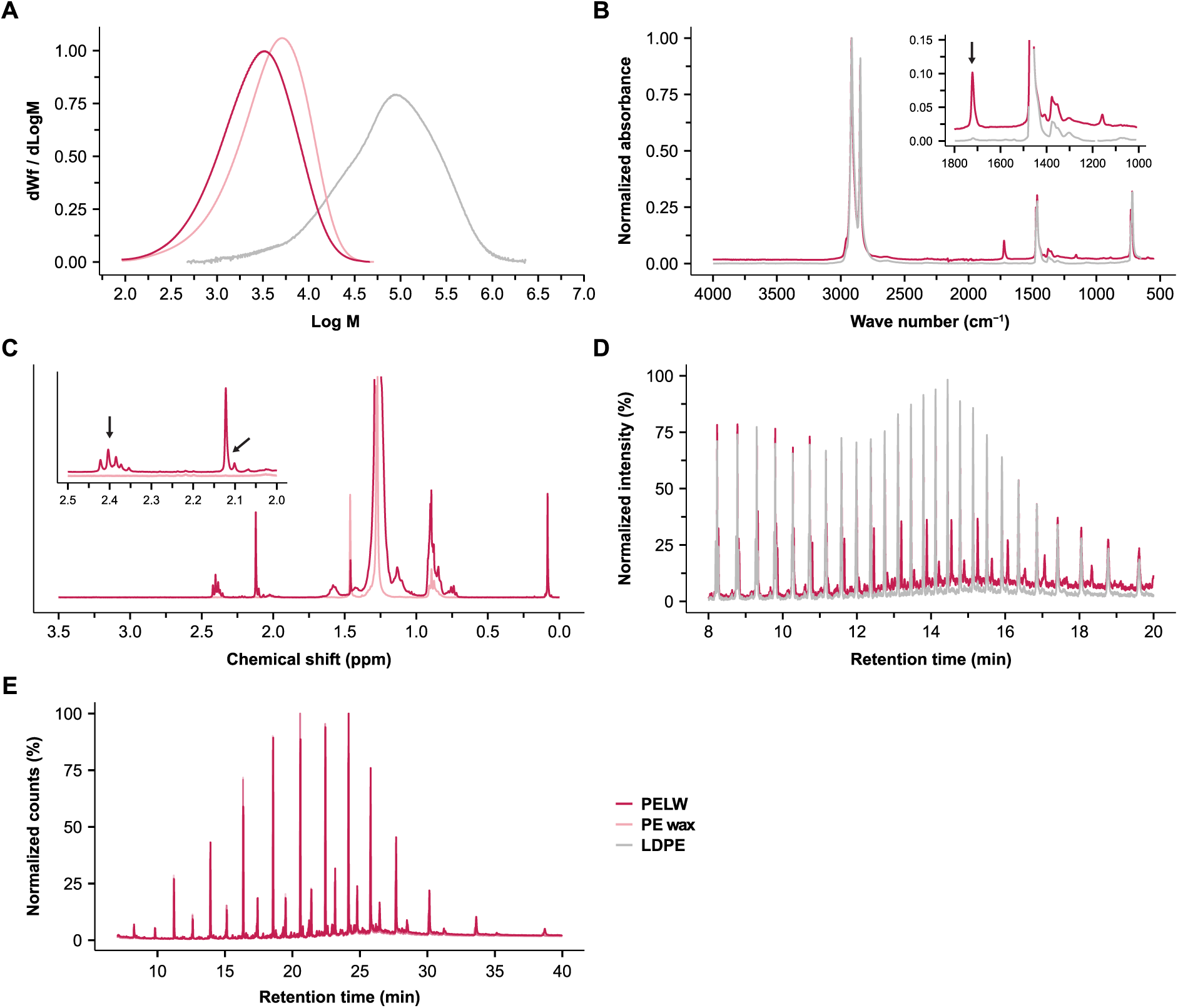
Characterization of PELW. Data for reference LDPE was obtained from Stepnov et al. [56]. **(A)** SEC analysis of PELW, a reference LDPE and a PE wax. The molar mass distribution of PELW is different than that of the PE wax and LDPE. **(B)** Overlayed FTIR spectra of the PELW and a reference LDPE. Compared to LDPE, the strong additional peak at ≈1722 cm^-1^ clearly indicates oxygen functionalities in the PELW (i.e. carbonyl groups in ketones or aldehydes). **(C)** NMR spectra of PELW. The singlets at 2.10-2.12 ppm and the triplet between 2.35-2.42 ppm confirm the presence of a carbonyl at the second carbon. **(D)** Overlaid Py-GC-MS chromatograms of PELW and LDPE. In the PELW, additional peaks were observed that were absent in the LDPE (see Table S2D for peak annotation). **(E)** GC-MS chromatograms of PELW ethyl acetate extracts, showing alkanes and 2-ketones between 10-35 carbons in chain length (see Table S2E for peak annotations). Reference LDPE data for panel **A** and **B** were obtained from Stepnov et al. [56]. Source data can be found in Table S2A-E.

The small singlet at 2.10 ppm showed a correlation to a carbonyl at 212.5 ppm, which stems from a 2-ketone next to a branching point. At 9.77 ppm, the ^1^H-NMR-spectrum also showed a triplet just above noise level, indicating that trace amounts of aldehydes are also present. The NMR data also provides means to quantify the degree of oxidation. Based on integrals in the ^1^H-NMR-spectrum, approximately 6‰ of all carbon atoms in the PE are oxidized to ketones. Of these ketones, 84% are 2-ketones. The Py-GC-MS data revealed the presence of analytes in the PELW sample that were not observed in the LDPE control sample (**Fig. 3D**). These peaks were identified as 2-ketones between 12-32 carbons, using a reference library (**Table S2D**). GC-MS analysis confirmed the presence of these small molecules by identifying hydrocarbons, including alkanes with 13-35 carbons and 2-ketones with 10-34 carbons in the PELW samples (**Fig. 3E, Table S2E**). The presence of short 2-ketones in the PELW introduces oxygen functionality, potentially increasing the bioavailability of the plastic compared to pristine LDPE.

### An *A. guilloiae* FS11 isolate catabolizes low molecular mass alkanes and 2-ketones present in PELW

To gain deeper insight into the bacterial metabolism of PELW constituents, we isolated the dominant species from the metagenomic analysis, namely *A. guillouiae* FS11. While not growing on pristine LDPE (**Fig. 4A**), the growth of *A. guillouiae* FS11 on PELW was remarkably rapid for an insoluble substrate, showing an increase in OD600 from 0.00 to 0.56 after 75 h incubation (**Fig. S4**). SEC analysis of the PELW particles after bacterial growth showed a reduction of the amount of alkanes smaller than ∼316 g/mol (**Fig. 4B**; see the zoomed-in view). Remarkably, the molecular mass distribution curve obtained for PELW after bacterial grow closely overlapped with that of the negative control sample, providing no evidence for degradation of polymeric PE substrate. GC-MS quantification of the PELW low molecular mass oligomers showed depletion of the smallest alkanes and 2-ketones and substantial reduction in medium length oligomers (10 to 35 carbons in length), compared to the control samples (**Fig. 4C**). Interestingly, the PELW particles that only were exposed to MM medium, also showed a partial reduction in short alkane and 2-ketone oligomers (**Fig. 4C**), suggesting formation of shorter derivatives that are likely easily accessible for microorganisms. Similar to the observations made for enrichment cultures growing on PELW (**Fig. 1C and D**), *A. guillouiae* FS11 formed biofilms on the surface of PELW particles (**Fig. 4D**).

**Fig. 4.**
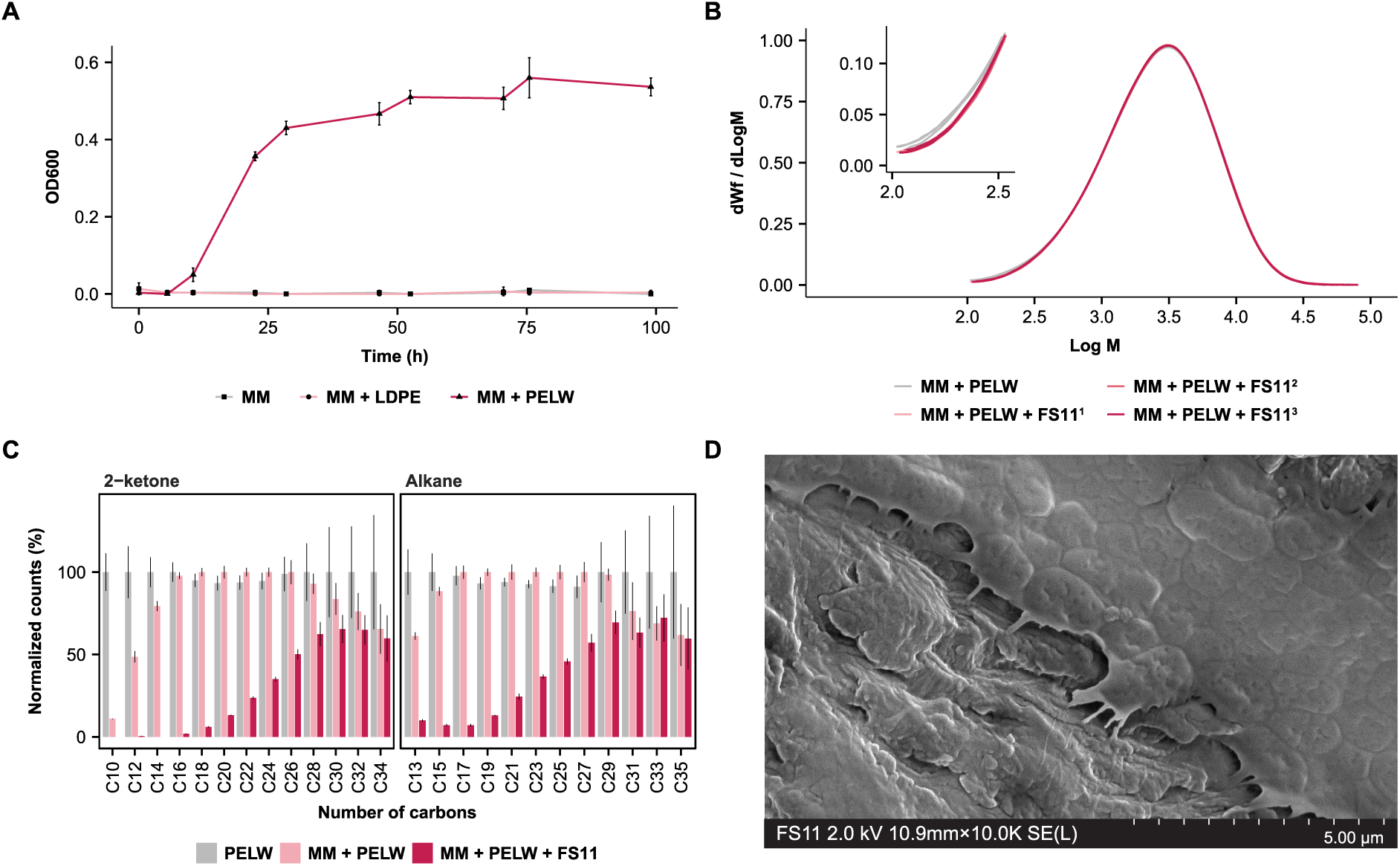
Growth profiles of *A. guillouiae* FS11 on PELW and subsequent characterization of the substrate. **(A)** Growth of A. guillouiae FS11 in minimal medium without any carbon source (MM) or supplemented with either PELW (MM+PELW) or LDPE (MM+LDPE). Data are averages ± standard deviations (error bars) of three biological replicates. **(B)** SEC chromatograms of PELW following incubation with MM or MM + *A. guillouiae* FS11. One control sample (MM + PELW) and three replicates (MM + PELW + FS11^1-3^) were set up. Sample-specific chromatograms are shown. Note that chromatograms are superimposing in the high molecular weight regions, indicating no degradation of polymeric fraction of PELW substrate, as the molar mass distribution would shift to the left. **(C)** GC-MS results of PELW showing the relative reduction of each compound after growth of *A. guillouiae* FS11. Compound loss is expressed relative to the amount of the same compound detected in PELW incubated in the absence of *A. guillouiae* FS11. **(D)** SEM image of *A. guillouiae* FS11 adhering to the surface of PELW. Source data for SEC and GC-MS can be found in Table S2F-G.

### *A. guillouiae* FS11 may use flavin-containing monooxygenases for 2-ketone conversion

To identify the enzymes involved in the degradation of the oligomeric alkanes and 2-ketones in PELW, we conducted a proteomic analysis of secreted, planktonic and biofilm-derived proteins produced by *A. guillouiae* FS11 during growth on PELW, triacontane and sodium succinate. Enzymes involved in alkane degradation, putative 2-ketone degradation and β-oxidation were generally more abundant under PELW conditions (**Fig. 5A**). In the biofilm-derived proteome, all known enzymes involved in alkane degradation, as well as the putative enzymes involved in 2-ketone oxidation were detected (**Fig. 5A**). Two LadA (LadA_3609, LadA_4534), were exclusively detected in the PELW-biofilm proteome. AlkB_3757 was detected in all proteomes, except for the succinate control. Finally, YiaY_433 and ALDH_4343 were among the most abundant proteins detected in the PELW-associated (biofilm, planktonic and secreted) and C30 (planktonic) proteomes (**Fig. 5A**), indicating that enzymes involved in the complete alkane degradation pathway are active under these conditions.

**Fig. 5.**
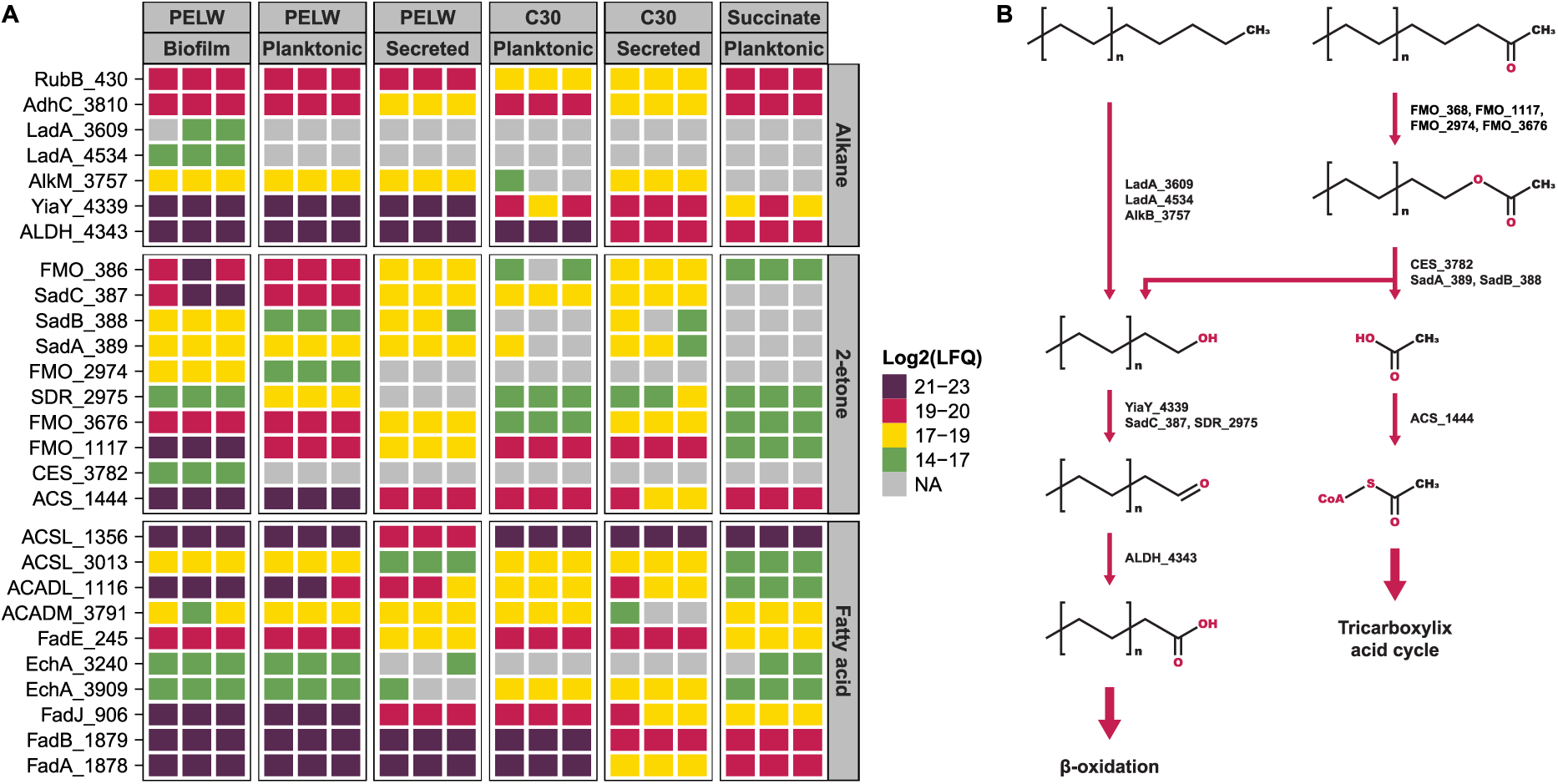
Selected differentially abundant proteins in the *A. guillouiae* FS11 proteome and proposed metabolic pathway for degradation of 2-ketones and alkanes in PELW. Panel A shows the LFQ intensities of the known and proposed enzymes in the pathways for terminal alkane oxidation, 2-ketone degradation and β-oxidation. Missing values are shown in grey. Panel B shows a proposed metabolic pathway of alkanes and 2-ketones found in PELW, with chain lengths between 10-35. Alkanes are terminally oxidized, while 2-ketones are oxidized by FMOs, generating esters. Esters are hydrolyzed into alcohols and acetate, which can enter the alkane degradation pathway and the tricarboxylic acid (TCA) cycle, respectively.

Several differentially abundant FMOs, that may be actively transforming 2-ketones, were identified in the PELW and C30 proteomes (**Fig. 5A**). These included the FMO_386, encoded by a gene associated with a cluster that was confirmed via BLASTn analysis to correspond to the *sad* gene cluster of *Acinetobacter sp.* strain NyZ410 [51]. FoldSeek predictions using AlphaFold3 models indicated that the FMO corresponds to SadD, a BVMO known to convert aliphatic ketones into esters [51]. Additional FMOs were detected at high abundance either exclusively in the PELW proteome (FMO_2974), or in the PELW or C30 proteome (FMO_3676, FMO_1117). All these FMOs exhibit high structural and sequence similarity to characterized BVMOs (**Fig. S6, Table S1J**). The presence of multiple FMOs in the proteome may reflect the need to accommodate the 2-ketones of different chain lengths detected in the PELW (**Fig. 4C**), as such protein redundancy is commonly considered a strategy to expand substrate range [52]. The esterases SadA_389 and SadB_388, together with the carboxylesterase CES_3782, exhibited significantly higher abundance in the proteomes derived from PELW and C30 compared to the control conditions (**Fig. 5A**). The hydrolysis products (an alcohol and acetate) derived from the activity of these enzymes can either enter the terminal alkane oxidation pathway or be converted into acetyl-CoA by the Acetyl-CoA Synthetase ACS_1444, respectively (**Fig. 5A**).

Collectively, the detection of the enzymes described above (**Fig. 5A**), in combination with the depletion of both alkanes and 2-ketones (**Fig. 4C**), indicate that the small molecules derived from the PELW can be consumed through terminal alkane oxidation and 2-ketone degradation, which is a shorter version of the subterminal oxidation of alkanes (**Fig. 5B**).

## Discussion

Biotechnological conversion of PE to useful products like platform chemicals or microbial single cell proteins is an important research area given the unfathomable amounts of PE waste available and continuously accumulating on the planet. Indeed, many studies exist where PE is reported to be microbially or enzymatically degraded [53,54], but few of these have been reproduced by others and some attempts of reproduction have failed [55]. Recently, several cautionary articles and reviews have been published that urge research to be more careful in experimental design and interpretation of results [18,55,56]. One common misunderstanding arises from the use of model PE substrates that do not have the properties of a typical industrial-grade PE. A very good example is the PELW used in the present study, which is marketed as “PE” by the supplier (Sigma-Aldrich). This product has been used in multitude of studies [15,43–50], many of which report microbial PE-degradation. From a polymer chemist standpoint, the Sigma-Aldrich PE is a PE-like wax due to its low M_w_ (∼3900 g/mol), and such a polymer cannot be compared to a typical commercial PE which has M_w_ often exceeding 100 000 g/mol, where the high molecular weight is what provides the high chemical and structural robustness. Our data clearly shows that bacteria like *A. guillouiae* F11, as well as microbial communities can efficiently utilize PELW as a sole carbon source by metabolizing the alkane oligomers and 2-ketones present in the wax (**Fig. 4 and Fig. S2**). In our commercial LDPE sample, no such oligomers were present (**Fig. 3**), thus neither the *A. guillouiae* F11 isolate, nor the soil derived microbial communities were able to grow (**Fig. S2 and Fig. S4**). It is not unlikely that the mechanism of PELW utilization in other studies, where the PELW is mistaken for a compound that has the properties of high-molecular mass PE (even referred to as LDPE at times, [43,45,47,48]), is the same as observed in our study. A second key observation made in the present study is the abundant presence of 2-keto groups in the PELW chains (**Fig. 3**), a feature absent from the supplier’s product specification, that could increase the material’s biodegradability. Interestingly, we are not the first to identify oxidation of the PELW. Zampolli et al. also performed GC-MS on the material (most likely a different batch) and reported the presence of multiple different oxygen-containing functional, groups, including 2-ketones [49]. These observations suggest that it is not only the product purchased for the present study that is oxidized, but possibly a larger batch of the product. Notably, FTIR analysis of two independently purchased PELW samples (batch numbers MKCT5408 and MKCP961) confirmed the same oxidation pattern (results not shown). Carbonyl groups can be introduced into PE, including the low molecular weight fractions, through i.e. chemical [57], thermal [58] and photo-oxidation [59], supporting the idea that these components are oxidation derivatives from original PELW material.

The mechanism by which the oligomeric alkanes and 2-ketones are consumed can give valuable insight into how PE waste potentially can be enzymatically depolymerized, given that the material is pretreated to reduce molecular weight, e.g. by oxidation. When searching for microorganisms with the capability to depolymerize PE, or oxidized PE components and derivatives, it is reasonable to focus on natural communities that have been exposed to plastic pollution over many years. In this work, an enrichment experiment using an oxidized PELW was established from soil communities from a landfill contaminated with weathered plastic waste. Shotgun metagenomic profiling showed that *Acinetobacter* and *Pseudomonas* spp. became abundant in the enrichment cultures on PELW, triacontane and tetracontane (**Fig. 1B**). Members of the genera *Pseudomonas* [60,61] and *Acinetobacter* [52,62] are some of the most studied alkane degrading microorganisms and have been proposed to be able to adhere and/or degrade PE [12,63–65].

Genome-centric metatranscriptomics showed that microbial communities enriched from soil collected at a plastic landfill could actively metabolize PELW as the sole carbon source, whereas no growth was observed in the pristine LDPE (**Fig. S2**). PELW contains shorter hydrocarbons, alkanes and 2-ketones (**Fig. 3**), which serve as substrates for the microbes [66]. Genes coding for enzymes for terminal oxidation of alkane, including the monooxygenases LadA and AlkB, were upregulated in the S1PELW, S1C30, S2C30 and S2C40 enrichments (**Fig. 2**). In the S2PELW and S1C40 enrichments, upregulated CYP genes may allow subterminal hydroxylation of alkanes (**Fig. 2C-D**), enabling the microorganisms to degrade alkanes without AlkB or LadA [42]. Notably, several genes encoding FMOs were upregulated on PELW, C30 and C40 (**Fig. 2**). The FMO family comprises a diverse range of xenobiotic-metabolizing enzymes, which use NADPH as a cofactor, and FAD as a prosthetic group. This family includes BVMOs that can also play a role in the subterminal oxidation of alkanes [67]. Here, an alkane monooxygenase introduces a hydroxyl group at a subterminal position of the alkane chain, which is further oxidized to a ketone by an alcohol dehydrogenase. The carbonyl group in the ketone can then converted into an ester group by a BVMO [66]. If these FMOs have BVMO functionality, they can contribute to the conversion of 2-ketones, derived from the PELW, or subterminally hydroxylated alkanes, into esters.

Following isolation of the dominant species involved in PELW degradation, *A. guillouiae* FS11, proteomics and substrate analyses further demonstrated that it metabolizes the smaller compounds (alkanes and 2-ketones) in the PELW but is not able to cleave the polymeric backbone (**Fig. 4**). Proteomic results revealed that *A. guillouiae* FS11 metabolizes PELW components via terminal alkane oxidation, β-oxidation, and the TCA cycle, with abundant AlkB, LadA, Adh, YiaY, ALDH, and several FMOs, including FMO_2974, uniquely detected in PELW-derived proteomes (**Fig. 5A**). FMOs may actively interact with the 2-ketones and alkanes present in the oxidized PELW via the subterminal oxidation pathway (**Fig. 5B**). The exact role of the putative BVMOs remains to be elucidated through biochemical characterization on these substrates.

In summary, our study provides a valuable approach for studying how microbial communities interact with PE and PE-derivatives, and results support two main conclusions. First, advanced substrate characterization before and after microbial growth is essential to convincingly support assumptions of microbes being able to break down the PE backbone and subsequent mineralization. Second, the soil microbial communities and bacterial isolate could efficiently utilize shorter-chain and oxidized-derivatives of PE, while evidence of degradation of longer chain, pristine LDPE was not obtained. These results, in combination with the lack of biochemically characterized enzymes which act on polymeric PE [68], indicate that efforts should be made to create new PE-like materials with oxygen functionality to add enzymatic recyclability. Even though PE in nature remains challenging, our study can help guide the field of microbial plastic degradation by highlighting the challenges associated with data interpretation, and to advise caution in future studies.

## Acknowledgements

This research was funded by the Research Council of Norway under grant agreement no. 326975 (Enzyclic project). Mass spectrometry-based proteomic analyses were conducted at the MS and Proteomics Core Facility, Norwegian University of Life Sciences. This facility is a member of the National Network of Advanced Proteomics Infrastructure (NAPI), which is funded by the Research Council of Norway INFRASTRUKTUR-program (project number: 295910). The authors acknowledge Elixir Norway, supported by the Research Council of Norway’s (NFR) grant agreement no. 322392, for the data deposition support received for this study. RRC would like to thank colleagues Asbjørn Iveland, Bavan Mylvaganam, Sara Rund Herum and Steffen Annfinsen for their valuable analytical support.

